# A pathologically expanded, clonal lineage of IL-21 producing CD4+ T cells drives Inflammatory neuropathy

**DOI:** 10.1101/2024.01.07.574553

**Authors:** Maryamsadat Seyedsadr, Madison Bang, Ethan McCarthy, Shirley Zhang, Ho-Chung Chen, Mahnia Mohebbi, Willy Hugo, Jason K. Whitmire, Melissa G. Lechner, Maureen A. Su

## Abstract

Inflammatory neuropathies, which include CIDP (chronic inflammatory demyelinating polyneuropathy) and GBS (Guillain Barre Syndrome), result from autoimmune destruction of the peripheral nervous system (PNS) and are characterized by progressive weakness and sensory loss. CD4+ T cells play a key role in the autoimmune destruction of the PNS. Yet, key properties of pathogenic CD4+ T cells remain incompletely understood. Here, we use paired scRNAseq and scTCRseq of peripheral nerves from an inflammatory neuropathy mouse model to identify IL-21 expressing CD4+ T cells that are clonally expanded and multifunctional. These IL-21-expressing CD4+ T cells are comprised of two transcriptionally distinct expanded populations, which express genes associated with Tfh and Tph subsets. Remarkably, TCR clonotypes are shared between these two IL-21-expressing populations, suggesting a common lineage differentiation pathway. Finally, we demonstrate that IL-21 signaling is required for neuropathy development and pathogenic T cell infiltration into peripheral nerves. IL-21 signaling upregulates CXCR6, a chemokine receptor that promotes CD4+ T cell localization in peripheral nerves. Together, these findings point to IL-21 signaling, Tfh/Tph differentiation, and CXCR6-mediated cellular localization as potential therapeutic targets in inflammatory neuropathies.

## Introduction

Inflammatory neuropathies, which include CIDP (chronic inflammatory demyelinating polyneuropathy) and GBS (Guillain Barre Syndrome) are characterized by debilitating weakness and sensory loss. Hallmarks of these conditions include autoimmune demyelination and immune cell infiltration of peripheral nerves^1,2^. IVIg (intravenous immunoglobulin) is a mainstay of therapy, but it fails to achieve significant clinical responses in one-third of CIDP patients^3^, and GBS is associated with a 6.6-fold increase in mortality even with IVIG therapy^4^. Moreover, IVIg has broad effects on the immune system that remain incompletely defined^5^. Despite the need for more effective, mechanism-based treatments, new therapeutic approaches have not been introduced since the 1990’s^6,7^. This contrasts with Multiple Sclerosis (MS), an autoimmune demyelinating condition of the central nervous system (CNS), in which >10 disease-modifying therapies have been FDA approved since 1994^8^.

Progress in developing new immunotherapeutic agents for inflammatory neuropathies has been hampered by a paucity of knowledge regarding fundamental aspects of autoimmune pathogenesis. A major breakthrough in addressing this need has been the recent development of mouse models of inflammatory neuropathies that recapitulate multiple aspects of human disease. For instance, NOD.Aire^GW/+^ mice develop spontaneous autoimmune peripheral polyneuropathy (SAPP) that is associated with demyelination and immune cell infiltration in peripheral nerves^9,10^. In this model, autoimmune-prone NOD mice harbor a partial loss-of-function G228W mutation in the *Aire* (*Autoimmune Regulator*) gene, which allows the escape of autoreactive T cells from thymic negative selection. The increased frequency of autoreactive T cells that recognize myelin-specific self-antigens predisposes to their activation, infiltration into peripheral nerves and destruction of myelin in peripheral nerves. Importantly, inflammatory neuropathy has also been reported in patients with mutations in the *AIRE* locus^11^, highlighting the importance of autoreactive T cells in driving autoimmune peripheral neuropathy across species.

Among T cells, CD4+ T cells in particular are critical in the pathogenesis of inflammatory neuropathies. CD4+ T cells are increased in peripheral nerves of patients with inflammatory neuropathies^12–14^ and SAPP mouse models^10^, suggesting a role for CD4+ T cells in peripheral nerve myelin destruction. Moreover, CD4+ T cells from neuropathic mice are sufficient to transfer SAPP to immunodeficient recipients^9,10^, and a myelin-specific CD4+ TCR transgenic mouse model spontaneously develops autoimmune peripheral neuropathy, suggesting that CD4+ T cells are sufficient for the development of autoimmunity^10,15,16^. Despite these findings that support the importance of CD4+ T cells, key properties of pathogenic CD4+ T cells in inflammatory neuropathies remain incompletely understood.

Here, we show that, in peripheral nerve infiltrates of neuropathic NOD.Aire^GW/+^ mice, terminally-differentiated effector CD4+ T cells are clonally expanded and express IL-21. These IL-21-producing cells can be grouped into two transcriptionally distinct populations, which resemble T follicular helper (Tfh) and T peripheral helper (Tph) cells. Notably, TCR clonotypes are shared in these two subsets, supporting a common lineage for these two populations. Additionally, we demonstrate that IL-21 signaling is required for neuropathy development, and that IL-21 upregulates CXCR6, a chemokine which promotes CD4+ T cell localization within peripheral nerves. Together, these findings demonstrate a critical role for IL-21 in disease pathogenesis and reveal multiple new molecular targets for the treatment of autoimmune peripheral neuropathies.

## Results

### IL-21 expressing CD4+ T cells are pathologically expanded in peripheral nerve autoimmunity

Previous studies have demonstrated infiltrating CD4+ T cells in peripheral nerve biopsies from both inflammatory neuropathy patients^12,13^ and SAPP mouse models^10,15^, supporting an important role for CD4+ T cells in the development of PNS autoimmunity. To gain deeper insights into the phenotype of pathogenic CD4+ T cells, we analyzed sciatic nerve infiltrating CD4+ T cells from neuropathic NOD.Aire^GW/+^ mice **(Fig. 1A)**. Single cell RNA sequencing (scRNAseq) analysis of 4,017 CD4+ T cells revealed 6 distinct groups **(Fig. 1B)**. These included clusters differentially expressing stem-like progenitor genes^17–19^ (*Tcf7*, *Klf2*, and *S1pr1;* Cluster 0); the early lymphocyte activation marker *Cd69*^20,21^ (Cluster 1); regulatory T cell (Treg) genes (*Foxp3*, *Il2ra;* Cluster 4); and a mix of lineage-associated genes (Cluster 5) **(Supp. Fig 1A,B)**.

**Figure 1.**
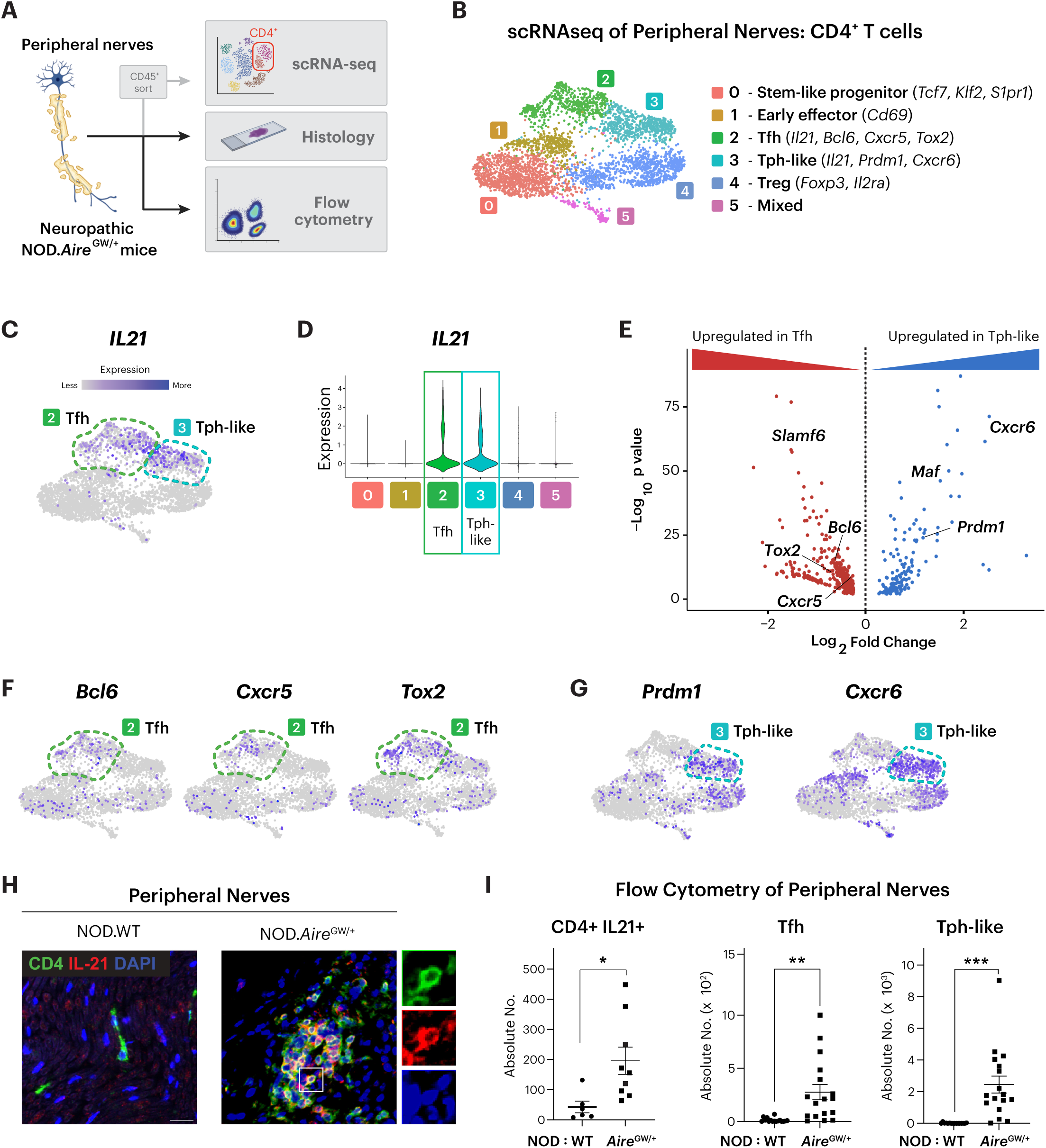
IL-21 expressing CD4+ T cells are pathologically increased in peripheral nerves of neuropathic NOD.Aire^GW/+^ mice. **(A)** Schematic for workflow to analyze peripheral nerve infiltrating CD4+ T cells in neuropathic NOD.Aire^GW/+^ mice. **(B)** UMAP plot of infiltrating CD4+ T cells in the sciatic nerves of NOD.Aire^GW/+^ mice (n=7). **(C and D)** Feature plot (C) and violin plot (D) of *Il21* expression. **(E)** Volcano plot of differentially expressed genes between cells in Tfh vs. Tph-like clusters (Log_2_FC>0.05, p_val_adj < 0.05). **(F-G)** Feature plots showing expression of key transcription factors and chemokines associated with Tfh (*Bcl6, Tox2, Cxcr5*) and Tph (*Prdm1, CXCR6*) cells. **(H)** Immunofluorescence staining and microscopy images of CD4, IL-21, and DAPI in NOD.WT (control) and neuropathic NOD.Aire^GW/+^ sciatic nerves. Right panels show the area outlined by white box, magnified, and split by fluorescence. **(I)** Absolute numbers of indicated cell types per sciatic nerve, as determined by flow cytometric analysis. CD4+ T cells that produce IL-21, Tfh cells (CD4+ ICOS+ PD1+ CXCR5+), and Tph-like cells (CD4+ ICOS+ PD1+ CXCR5-CXCR6+) were compared between NOD.WT vs. neuropathic NOD.Aire^GW/+^ sciatic nerves. Mann Whitney test; *p<0.05; **p<0.01; ***p<0.001.

Remarkably, two CD4+ T cell populations (Clusters 2 and 3) significantly upregulated IL-21 **(Fig. 1C,D; Supp. Fig. 1A),** a cytokine linked to Type 1 Diabetes (T1D) and other autoimmune conditions but not yet to PNS autoimmunity^22,23^. In contrast, IL-21 expression was absent in immune cells found in sciatic nerves of non-neuropathic NOD wildtype (NOD.WT) mice **(Supp. Fig. 1C,D)**^24^. Thus, the development of autoimmune peripheral neuropathy in NOD.Aire^GW/+^ mice is associated with IL-21 upregulation in peripheral nerve CD4+ T cells.

Comparison of the two IL-21 producing populations (Clusters 2 and 3) showed distinct transcriptional profiles, with 560 differentially regulated genes (p value adjusted<0.05; log_2_FC>0.5). Expression of genes (*Bcl6, Tox2, Cxcr5, Slamf6)* associated with Tfh cells^25,26^ were significantly upregulated in Cluster 2, while expression of genes *[Prdm1* (Blimp1), *Cxcr6]* associated with Tph cells^27–29^ were significantly upregulated in Cluster 3 (**Fig. 1E**). In accord, feature plots showed Tfh-associated genes (*Bcl6, Tox2*, *Cxcr5)* expressed by cells in Cluster 2 and Tph-associated genes (*Prdm1* and *Cxcr6)* genes expressed by cells in Cluster 3 (**Fig. 1F,G)**. Of note, Cluster 3 failed to express a subset of Tph-associated chemokines (e.g., *Cxcl13, Ccr2, Cx3cr1*) described in rheumatoid arthritis and other conditions^27^, a finding consistent with previous reports that chemokine expression by Tph cells is context dependent^30^. Together, these data suggest that Tfh (Cluster 2) and Tph-like (Cluster 3) cells are the primary source of IL-21 within the inflamed nerve.

We verified these scRNAseq findings using multiple complementary approaches. First, immunofluorescence staining of frozen sciatic nerve sections revealed co-localization of CD4 (green) and IL-21 (red) in neuropathic NOD.Aire^GW/+^ sciatic nerves, supporting the production of IL-21 by CD4+ T cells in inflamed peripheral nerves. Moreover, CD4 and IL-21 staining were absent in non-neuropathic controls (NOD.WT) **(Fig. 1H)**, which suggests that neuropathy is associated with increased IL-21 producing CD4+ T cells in peripheral nerves. Second, flow cytometric analysis of intracellular IL-21 cytokine staining showed accumulation of IL-21+ CD4+ T cells in NOD.Aire^GW/+^ sciatic nerves but not in non-neuropathic NOD.WT controls **(Supp. Fig. 2A; Fig. 1I)**. Collectively, these data support our scRNAseq analysis that IL-21 producing CD4+ T cells are pathologically expanded within the sciatic nerves of neuropathic NOD.Aire^GW/+^ mice.

Additionally, we used flow cytometry to quantify Tfh and Tph-like cells in inflamed nerves. Tfh cells are classically identified as CD4+ ICOS+ PD1+ CXCR5+ by flow cytometry, while Tph cells lack CXCR5 expression and are therefore identified as CD4+ ICOS+ PD1+ CXCR5-^25,26,30^ **(Supp. Fig. 2B)**. Because our scRNAseq analysis showed CXCR6 upregulation in the cluster of Tph-like cells **(Fig. 1E,G; Supp. Fig. 1A)**, we additionally incorporated CXCR6 as a marker for Tph-like cells **(Supp. Fig. 2B)**. Within the immune cell infiltrate of sciatic nerves of neuropathic NOD.Aire^GW/+^ mice, we found increased numbers of Tfh and Tph cells compared to non-neuropathic NOD.WT controls **(Fig. 1I)**. These findings, together with our scRNAseq analyses, reveal pathologic expansion of IL-21 producing Tfh and Tph-like populations in inflamed nerves of neuropathic mice.

### IL-21 producing cells in infiltrated peripheral nerves share a common lineage progenitor

Tfh and Tph cells are reported to share a number of phenotypic features, including IL-21 cytokine expression, presence in inflamed tissue; function in promoting B and T cell activation and maturation; and expression of the cell surface proteins ICOS and PD1^27,30,31^. These findings suggest a close molecular relationship between these two populations. On the other hand, Tfh and Tph cells are reported to be transcriptionally distinct^27^, and we found that, in inflamed peripheral nerves, Tfh and Tph-like cells have divergent gene expression profiles (**Fig. 1E).** Thus, it remains unclear whether Tfh and Tph-like cells are developmentally related, both in inflamed nerves and other contexts.

To begin to define the ontogeny of these cells, we examined transcriptional transitions of PNS-infiltrating CD4+ T cells from neuropathic NOD.Aire^GW/+^ mice. Slingshot pseudotime analysis of our scRNAseq data revealed three trajectories **(Supp. Fig. 2C)**, including one trajectory in which CD4+ T cells originated from stem-like progenitors (*Tcf7*+ *S1pr1*+ *Klf2+*), progressed through early effectors (*Cd69*+) and Tfh cell (*Bcl6+ Tox2+ Cxcr5+*) states, and terminated with Tph-like cells (*Prdm1*+ *Cxcr6+ Cxcr5-*) **(Fig. 2A)**. As cells progressed from stem-like progenitors, they decreased expression of stem-like progenitor transcription factor *Tcf7*, while upregulating the Tfh-associated transcription factor *Bcl6* and finally the Tph-associated transcription factor *Prdm1* **(Fig. 2B)**. These data suggest that Tph-like cells in SAPP arise from the Tfh population. Moreover, CD4+ T cells not only upregulated *Il21* as they progressed along this trajectory, but also *Ifng* and *Il10*. Because IFN-γ and IL-10 have both been identified as disease-promoting cytokines in SAPP^32,33^, these findings suggest that CD4+ T cells acquire an autoimmune effector phenotype as they differentiate along this lineage toward Tph-like cells.

**Fig. 2.**
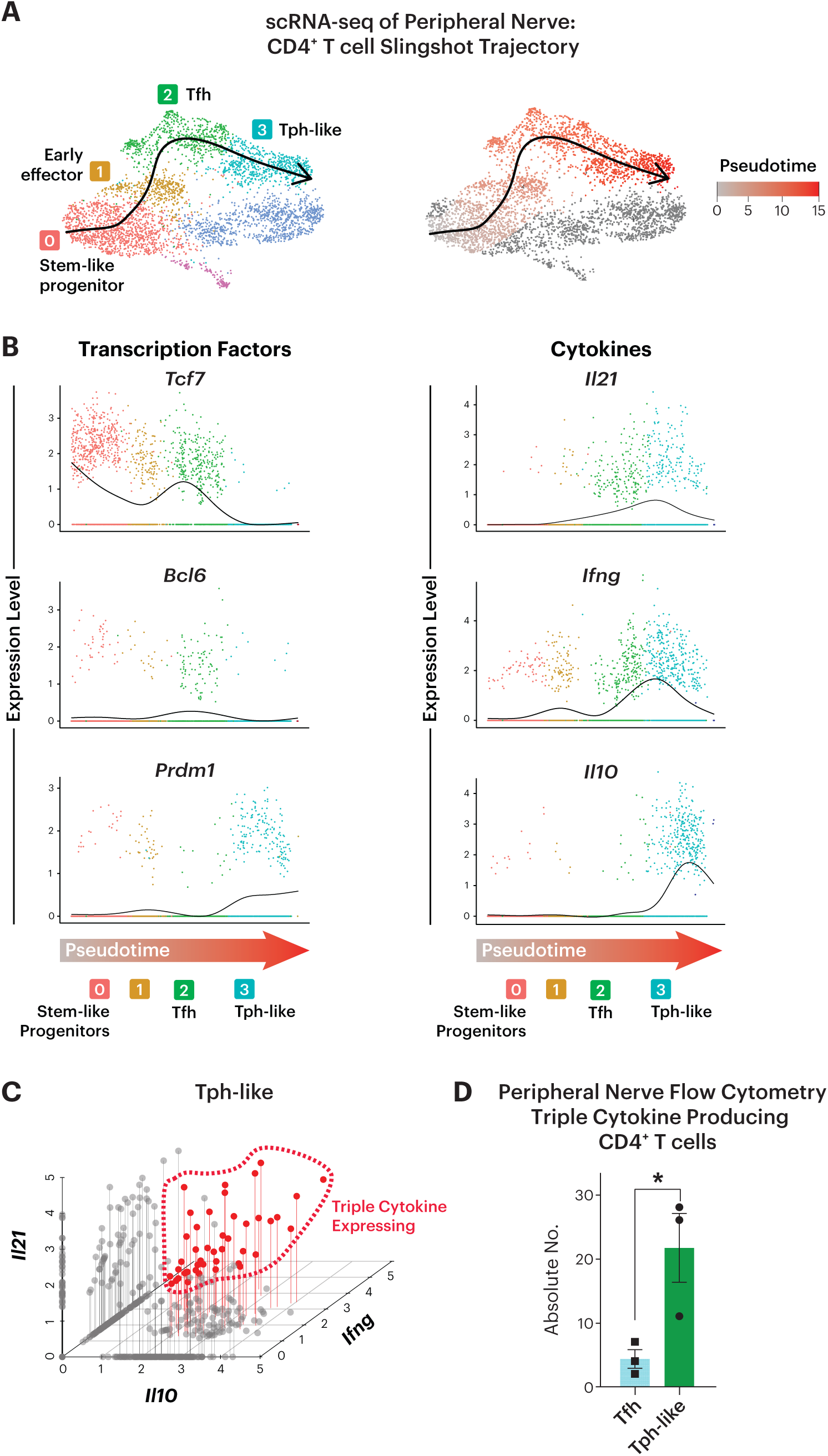
IL-21 producing cells in infiltrated peripheral nerves share a common lineage. **(A)** UMAP plot of CD4+ T cells with overlaid Slingshot pseudotime trajectory (left) and with cells color-coded chronologically along pseudotime, with gray representing the least differentiated and red indicating the most differentiated. **(B)** Expression of key genes along Slingshot pseudotime trajectory, color-coded by clusters shown in (B, left). **(C)** *Il21, Ifng,* and *Il10* co-expression by single cells within the Tph-like cluster. Triple cytokine-producing cells are circled. **(D)** Flow cytometric analysis of intracellular IL-21, IFN-y and Il-10 staining of peripheral nerve Tfh and Tph-like cells from neuropathic NOD.Aire^GW/+^peripheral nerves. Unpaired t-test; *p<0.05.

While our data show that IL-21, IFN-γ, and IL-10 expression are each highest in cells at the end of the pseudotime trajectory **(Fig. 2B)**, whether a single CD4+ T cell is capable of transcribing all three cytokines is unclear. To assess this, we correlated the expression of IL-21, IFN-γ and IL-10 within single cells. Using our scRNA-seq dataset to query cytokine expression in a single Tph-like cell, we found that 26% of cells expressed two of the three cytokines and 7% expressed all three **(Fig. 2C)**. The multifunctionality of CD4+ T cells was confirmed by intracellular cytokine staining and flow cytometric analysis (**Fig. 2D),** which demonstrated triple cytokine expression by a subset of Tph-like (CD4+ ICOS+ PD1+ CXCR5-) cells in neuropathic NOD.Aire^GW/+^ sciatic nerves. Thus, simultaneous expression of IL-21, IFN-γ, and IL-10 is seen in a subset of Tph-like cells in inflamed nerves of SAPP mice. This finding mirrors the co-expression of IL-21, IFN-γ, and IL-10 by pathogenic Tph cells in rheumatoid arthritis^27^ and Tph-like cells in kidney injury^34^.

### Multifunctional Tph-like cells are clonally expanded in peripheral nerves

During an autoimmune response, self-reactive T cells undergo clonal expansion with T cell receptor (TCR) self-antigen recognition and subsequent activation. Given the extremely low probability that somatic recombination at the TCR locus will result in the exact V(D)J rearrangement more than once, TCR sequences can be used as unique identifiers of T cell clones^35^. To query the clonality of PNS-infiltrating CD4+ T cells in neuropathic mice, we analyzed data from paired single cell TCR sequencing (scTCRseq) and scRNAseq of 4 NOD.Aire^GW/+^ sciatic nerve samples. The Treg and Mixed clusters were removed from this analysis in order to focus on conventional T (Tconv) cells. Using the total number of cells expressing each TCR sequence, we categorized each clonotype expansion as small (1<x<=5), medium (5<x<=20), or large (x>20). Most cells associated with medium and large clonal expansions mapped to the Tfh and Tph-like clusters **(Fig. 3A,B)**. Clonality was also measured using the Shannon entropy-based STARTRAC clonality index^36,37^, which demonstrated the highest Index Scores in Tfh and Tph-like populations **(Fig. 3C)**. Thus, the greatest degree of clonal expansion occurred in Tfh and Tph-like groups.

**Fig. 3.**
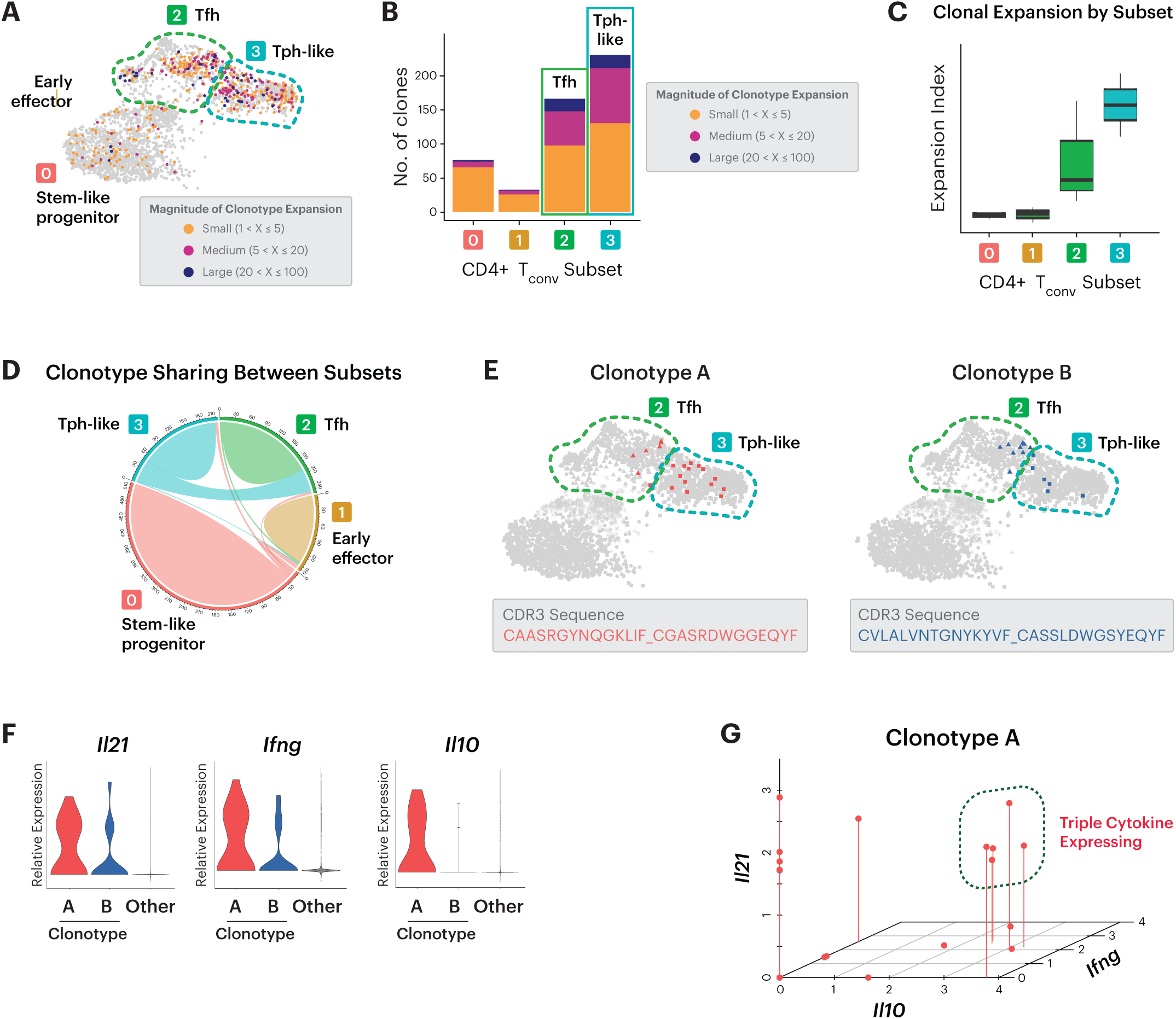
Tph-like cells in infiltrated peripheral nerves are clonally expanded and express IL-21, IFN-γ, and IL-10. UMAP of peripheral nerve-infiltrating CD4+ T conventional (Tconv) cells with projection of expanded clonotypes. Magnitude of expansion is grouped as small, medium, and large as indicated, with individual cells color-coded according to expansion magnitude. **(B)** Numbers of clonally expanded, Tconv cells, grouped by cluster. Degree of expansion indicated by color coding. **(C)** Clonal expansion levels of Tconv clusters quantified by STARTRAC. **(D)** Chord diagram of clonotype interconnections between clusters. Greatest sharing is seen between Tfh and Tph-like clusters (green). **(E)** Visualization of two expanded clonotypes (Clonotypes A and B) by their projection to the UMAP of CD4+ Tconv cells. CDR3 sequences for these two clonotypes are indicated. **(F)** Violin plots of cytokine expression levels in Clonotype A and B compared to all other cells. **(G)** Correlation plot showing levels of co-expression of *Ifng, Il10* and *Il21* cytokines by individual cells from Clonotype A. Cells expressing all three cytokines are circled.

Because of the low likelihood that two identical TCR sequences would arise independently in the same mouse, clonal sharing amongst cells with distinct phenotypes would suggests development from a common progenitor^35^. Visualization by chord diagram revealed the majority of clonal sharing occurred between Tfh and Tph-like clusters **(Fig. 3D)**^38,39^. Mapping of individual cells belonging to specific highly expanded clonotypes (i.e. Clonotype A and Clonotype B) also demonstrated sharing between Tfh and Tph-like cells **(Fig. 3E)**. For instance, cells from Clonotype A mapped to both Tfh and Tph-like clusters. This evidence for clonotype sharing between cells in the Tfh and Tph-like clusters suggests that Tfh and Tph-like cells originate from a shared precursor. These cells then proliferate in response to TCR activation and differentiate into distinct subsets.

We next examined the link between cytokine expression and clonal expansion **(Fig. 3F)**. Notably, *Il21* was highly expressed by cells associated with Clonotypes A and B, compared to all other cells. Similarly, *Ifng* was also highly expressed by Clonotypes A and B. *IL10*, however, was highly expressed by Clonotype A but not in Clonotype B. Interestingly, analysis of gene expression within single cells revealed a subset of cells associated with Clonotype A that simultaneously expressed all three cytokines **(Fig. 3G**). Together, these data identify clonally-expanded cells that traverse Tfh and Tph-like clusters and are capable of simultaneously expressing IL-21 and the pathogenic cytokines IFN-γ and IL-10.

### IL-21 signaling is essential for the development of autoimmune peripheral neuropathy

Although IFN-γ and IL-10 have been implicated in SAPP pathogenesis^32,33^, the role of IL-21 remains unclear. Upregulation of IL-21 in infiltrating CD4+ T cells and its expression by clonally expanded T cells suggests a critical role for IL-21 signaling in PNS autoimmunity development. To investigate this, we generated female NOD.Aire^GW/+^ mice with one or two copies of loss-of-function mutations in IL-21 receptor. Female NOD.Aire^GW/+^ mice with a heterozygous mutation in IL-21R (NOD.Aire^GW/+^ IL-21R^Het^ mice), developed neuropathy with the same onset and incidence as NOD.Aire^GW/+^ mice sufficient for IL-21R (NOD.Aire^GW/+^ IL21R^WT^) **(Fig. 4A)**. In contrast, female NOD.Aire^GW/+^ mice with homozygous mutations in IL-21R (NOD.Aire^GW/+^ IL21^KO^ mice) were protected against SAPP **(Fig. 4A). T**his protective effect of IL-21R deficiency was not sex-dependent, since Il-21R deficiency was also protective in male NOD.Aire^GW/+^ mice **(Supp. Fig. 3A).**

**Fig. 4.**
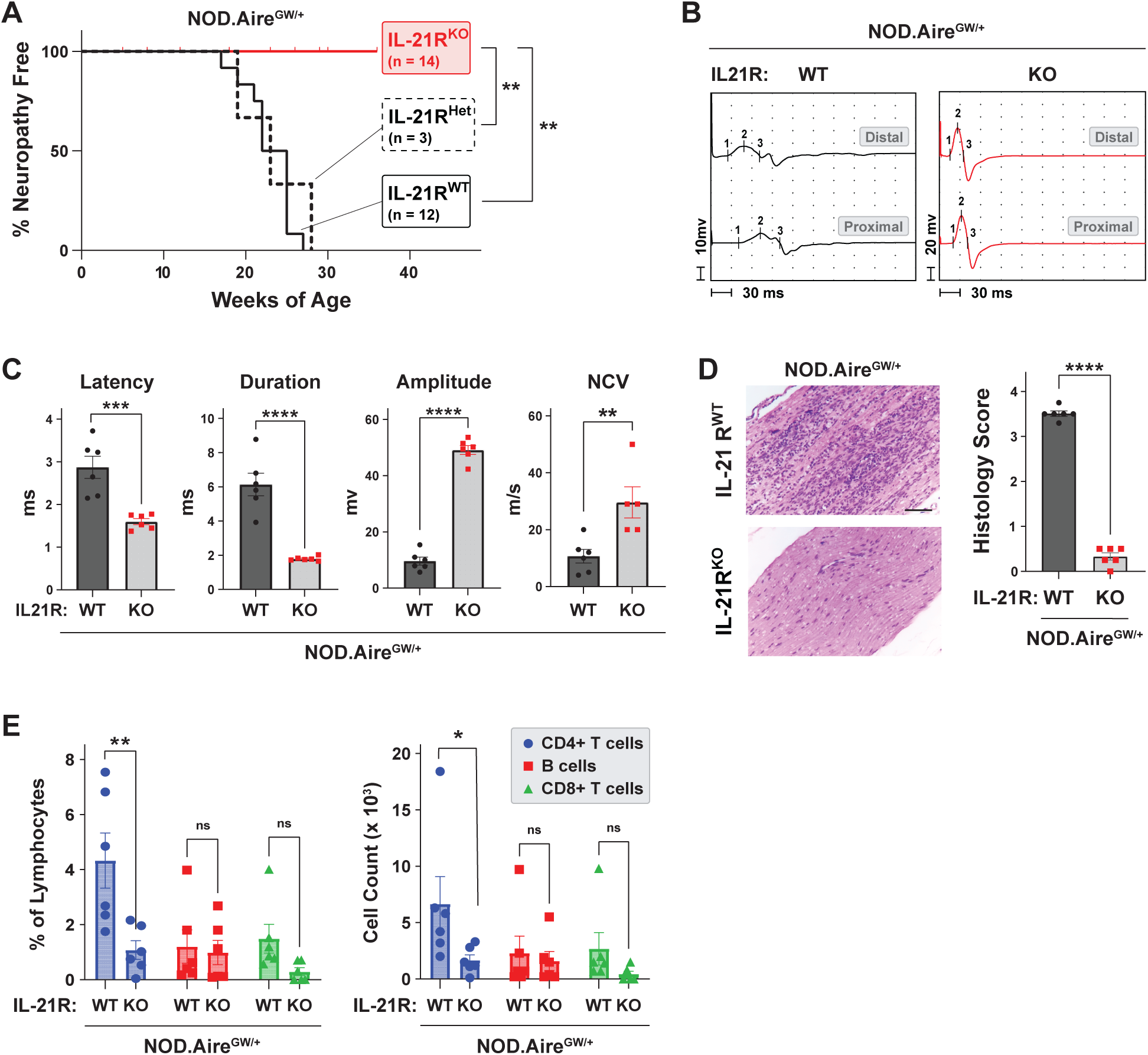
IL-21R signaling is required for neuropathy development in NOD.Aire^GW/+^ mice. **(A)** Neuropathy free incidence curve for female NOD.Aire^GW/+^ IL-21R^KO^ vs. NOD.Aire^GW/+^ IL-21R^Het^ vs. NOD.Aire^GW/+^ IL-21R^WT^ mice. Mantel-Cox test, **p<0.01. **(B,C)** Representative distal and proximal compound muscle action potentials from IL-21R-sufficient vs. deficient NOD.Aire^GW/+^ sciatic nerves. The latency, duration, amplitude, and nerve conduction velocity (NCV) were quantified and compared between groups (n=6, unpaired t-test). **(D)** The immune cells infiltration scores in the forelimb nerves assessed by H&E staining and compared between groups (n=6, unpaired t-test, scale bar=200uM). **(E)** The frequency and cell number of CD4+ T cells, B-lymphocytes (CD4-CD8-B220+), and CD8+ T cells compared between groups in the peripheral nerves (n=6, 2-way ANOVA). Bonferroni’s multiple correction; **p<0.01.

We have previously reported that female NOD.Aire^GW/+^ mice show evidence of demyelination on motor nerve electrophysiology^32^. In comparison, compound muscle action potentials from IL-21R deficient NOD.Aire^GW/+^ mice showed improvement in multiple parameters, including reduced latency and duration, and increased amplitude and nerve conduction velocity (NCV) **(Fig. 4B,C).** Additionally, histological analysis revealed significantly reduced peripheral nerve infiltration in NOD.Aire^GW/+^ IL-21R^KO^ mice, compared to NOD.Aire^GW/+^ IL-21R^WT^ **(Fig. 4D)**. Thus, our findings indicate a critical role for IL-21 signaling in the development of SAPP.

IL-21 has broad cellular targets, since IL-21R is expressed by various immune cell types (e.g., CD4+ T cells, CD8+ T cells, B cells)^40^. Flow cytometric analysis of peripheral nerve immune infiltrate indicated that decreased PNS immune infiltration in IL21R-deficient mice is due to lower numbers of CD4+ T cells, with no significant change in CD8+ T cells or B220+ B cells **(Fig. 4E).** Further, peripheral nerve infiltrating CD4+ T cells from IL21R-deficient mice demonstrated a decrease in IL-21, IFN-γ, and IL-10-producing CD4+ T cells **(Supp. Fig. 3B)**, suggesting that IL-21 from CD4+ T cells signals in an autocrine manner to increase numbers of cytokine-producing CD4+ T cells in inflamed peripheral nerves. However, it remains unclear the IL-21 regulated signals important for positioning pathogenic CD4+ T cells within inflamed peripheral nerves.

### CXCR6 upregulation in CD4+ T cells is IL-21-dependent

Our initial scRNAseq analysis revealed that the chemokine receptor CXCR6 is upregulated in Tph-like cells in peripheral nerves of neuropathic NOD.Aire^GW/+^ mice **(Fig. 1G; Supp. Fig. 1A)**. Notably, flow cytometric analysis of Tph-like (CD4+ PD1+ CXCR5-) cells in peripheral nerves revealed significantly lower CXCR6 mean fluorescence intensity (MFI) in IL-21R-deficient NOD.Aire^GW/+^ mice, compared to IL-21R-sufficient controls **(Fig. 5A).** These *in vivo* findings are in accord with previously published microarray data, which show that IL-21 stimulation *in vitro* upregulates CD4+ T cell expression of *Cxcr6* **(Supp. Fig. 3C)**^41^. Thus, CXCR6 expression by CD4+ T cells is IL-21-dependent.

**Fig. 5.**
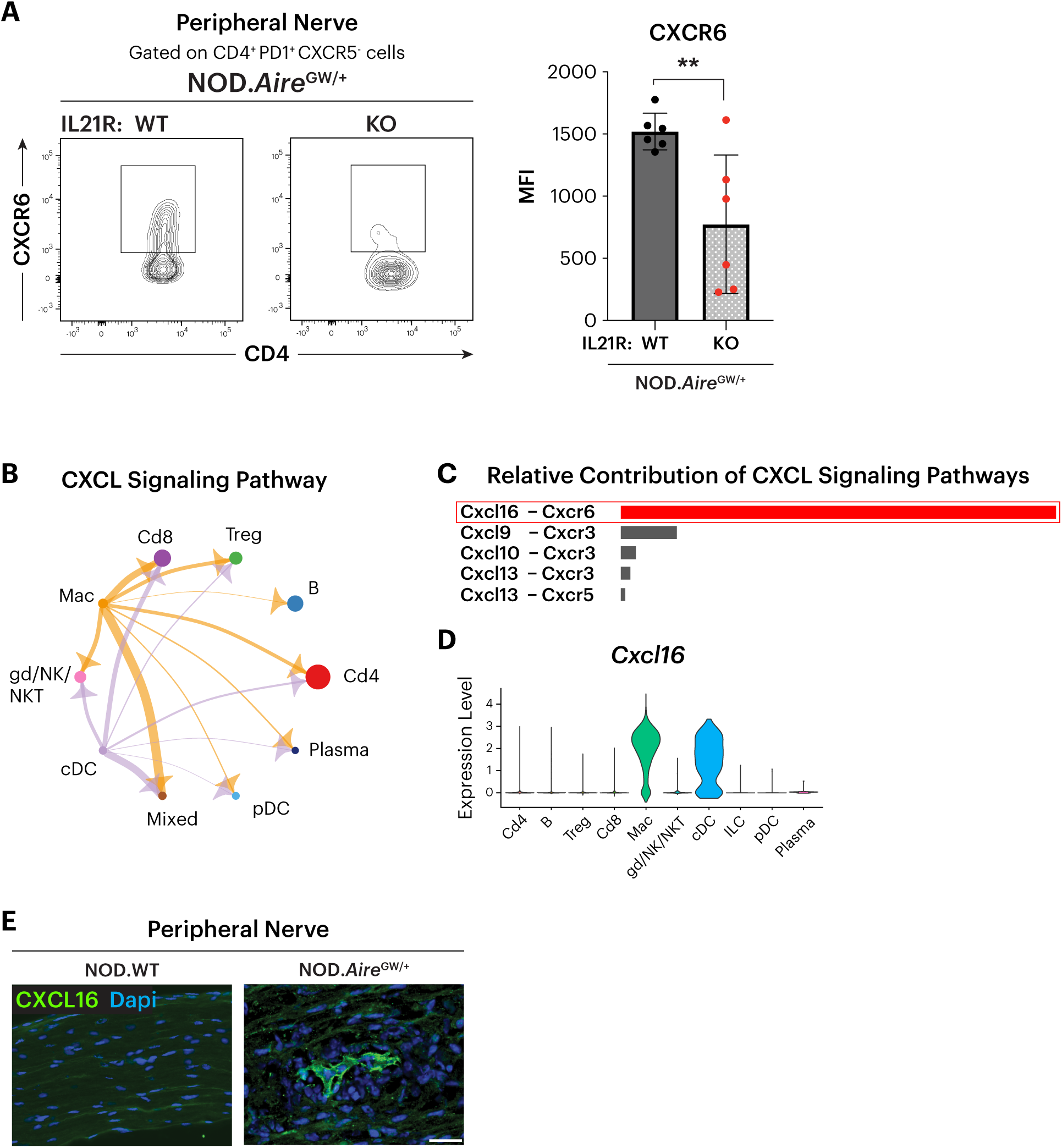
IL-21 mediated CXCR6 upregulation is required for the recruitment of CD4+ cells to the peripheral nerves. **(A)** Flow cytometric analysis of CXCR6 expression on CD4+ Tph cells from IL21R-sufficient vs. deficient NOD.Aire^GW/+^ sciatic nerves. **(B)** CellChat analysis of ligand-receptor interactions show upregulation of the ‘CXCL signaling pathway’. **(C)** The relative contribution of chemokine ligands and corresponding receptors in the sciatic nerves of NOD.Aire^GW/+^ mice. **(D)** Expression of CXCL16 in infiltrated immune cell subsets. Highest expression is seen in macrophages (Mac) and conventional dendritic cells (cDCs). **(E)** Immunostaining of CXCL16 in the peripheral nerves of a NOD.WT and NOD.Aire^GW/+^ neuropathic mice (scale bar=20µm).

To identify molecular mechanisms governing T cell positioning within inflamed peripheral nerves, we analyzed a previously published scRNAseq dataset of NOD.Aire^GW/+^ nerve-infiltrating immune cells (GSE180498). Using the CellChat R package to characterize ligand:receptor interactions, we identified significant upregulation of the ‘CXCL signaling pathway’ **(Fig. 5B)**, with prominent interactions between myeloid cells and T cells. Of these interactions, Cxcl16-Cxcr6 pairs were the most upregulated of the ‘CXCL signaling pathways’ **(Fig. 5C)**. Cxcl16 is the only known ligand for Cxcr6, and Cxcl16-Cxcr6 interactions have been reported to play an important role in positioning T cells in tumors and other tissues^42–44^. Whether Cxcl16-Cxcr6 interactions plays a role in positioning pathogenic T cells in inflamed peripheral nerves, however, is unknown.

In support of an important role for Cxcl16-Cxcr6 interactions, *Cxcl16* and *Cxcr6* expression are higher in infiltrating immune cells of neuropathic NOD.Aire^GW/+^ nerves, compared to non-neuropathic NOD.WT controls **(Supp. Fig. 4A)**. *Cxcl16* is highly expressed by macrophages and conventional dendritic cells (cDCs) in neuropathic NOD.Aire^GW/+^ nerves, while *Cxcr6* is expressed by lymphocytes **(Fig. 5D, Supp. Fig. 4A)**. Immunofluorescence staining of peripheral nerves from NOD.Aire^GW/+^ mice confirmed CXCL16 expression, which is absent in non-neuropathic NOD.WT controls **(Fig. 5E)**. Together, these data led us to hypothesize that Cxcl16-Cxcr6 interactions are important in positioning CD4+ T cells within inflamed peripheral nerves, and that downregulation of Cxcr6 with IL-21R deficiency prevents CD4+ T cell accumulation in inflamed peripheral nerves.

### IL-21-dependent CXCR6 assists autoreactive CD4+ T cell localization to the peripheral nerve

To test the role of Cxcl16-Cxcr6 interactions, we transduced neuropathic NOD.Aire^GW/+^ splenic CD4+ T cells with a viral vector co-expressing Cxcr6 and a GFP reporter **(Fig. 6A)**. As a negative control, cells were transduced with an empty vector expressing only an mCherry reporter. To determine the *in vivo* capacity of Cxcr6-overexpressing CD4+ T cells to localize to peripheral nerves, Cxcr6-overexpressing and control cells were sorted by reporter gene expression and co-transferred as a 1:1 mix to the same NOD.SCID recipient **(Fig. 6B).** This allowed for assessment of both CD4+ T cell groups within the same host environment. 4-5 weeks after adoptive transfer, cell distribution was assessed by flow cytometry. While the relative numbers of CXCR6-overexpressing cells to control (GFP : mCherry) cells were approximately equivalent in spleen and lymph nodes, the relative numbers of CXCR6-overexpressing cells were higher in the peripheral nerves **(Fig. 6C**). In parallel, we also transferred sorted CXCR6-overexpressing and control cells into separate NOD.SCID hosts **(Fig. 6D)**. In this experimental setup, the absolute number of CXCR6-overexpressing CD4+ T cells was also increased in peripheral nerves, compared to control CD4+ T cells **(Fig. 6E).** In contrast, no differences in CD4+ cell counts were observed in the spleen or lymph nodes. Together, these data support a model in which IL-21-dependent expression of CXCR6 in CD4+ T cells promotes their localization within the inflamed tissues of the peripheral nervous system **(Fig. 6F)**.

**Figure 6.**
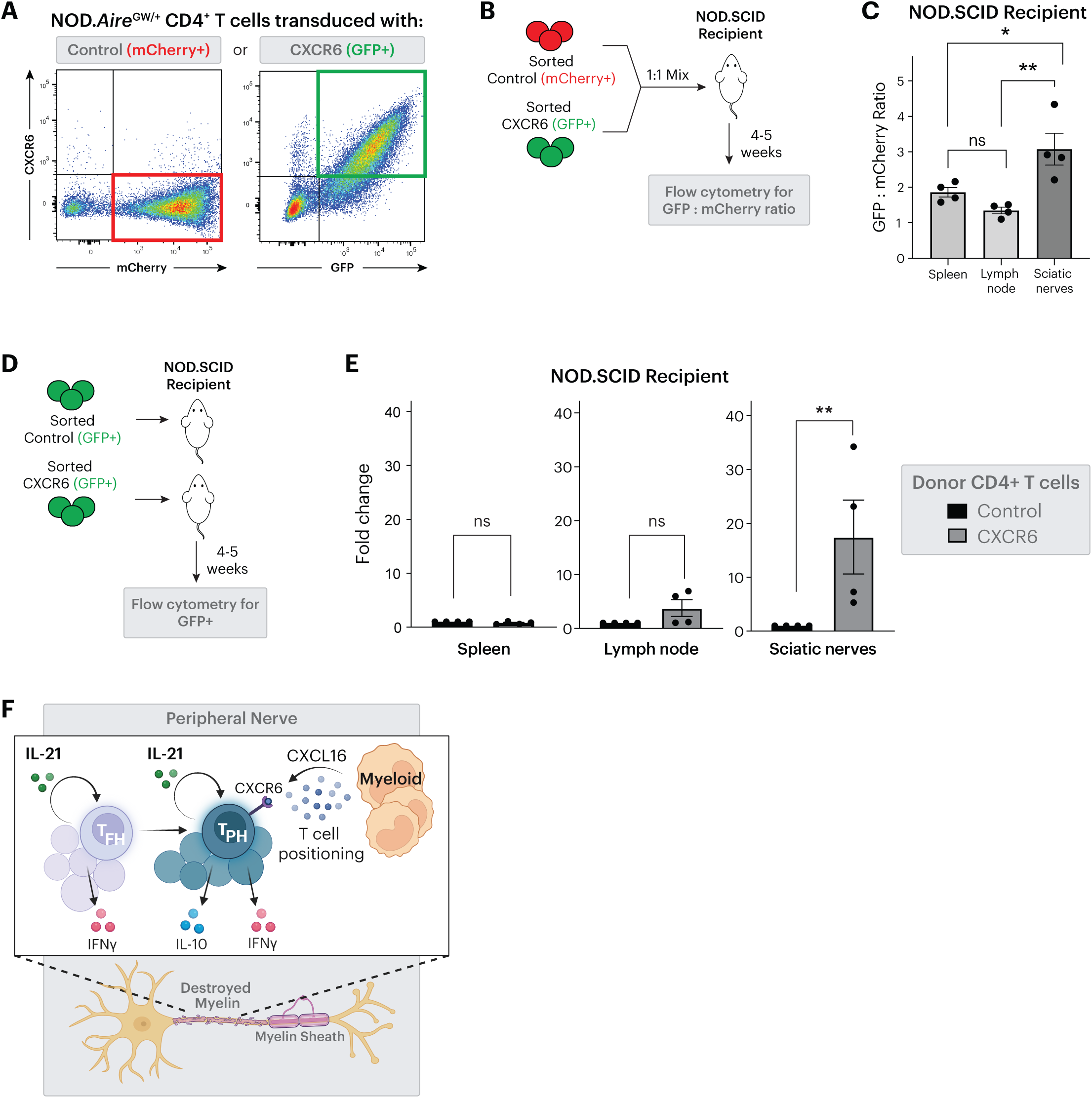
CXCR6 facilitates CD4+ T cell localization to inflamed peripheral nerves. **(A)** Flow cytometry plots of CD4+ cells from NOD.Aire^GW/+^ mice transduced with empty vector (mCherry) or CXCR6-GFP vector. **(B)** Experimental design for adoptive transfer of CXCR6 and control vector transduced CD4+ cells, mixed 1:1 prior to transfer into NOD.SCID recipients. **(C)** Quantification of GFP (CXCR6 vector) to mCherry (empty vector) ratio in spleen, lymph node, and sciatic nerve of recipient SCID mice (n=4, paired t test). **(D)** Experimental design for adoptive transfer of CXCR6 or control vector transduced CD4+ cells into NOD.SCID recipients. **(E)** The fold change difference in sciatic nerve, lymph nodes, and spleen calculated by normalizing the number of cells to the control transduced group (n=5, paired t-test). **(F)** Model for how pathogenic CD4+ T cells accumulate in inflamed peripheral nerves. Autocrine IL-21 signaling leads to Cxcr6 upregulation in Tph-like cells, which allows Cxcr6 to interact with Cxcl16 expressed by myeloid cells in inflamed peripheral nerves.

## Discussion

Understanding the autoimmune pathogenesis of inflammatory neuropathies has been greatly facilitated by the development of SAPP mouse models. These models, along with clinical observations in patients, have demonstrated a critical role for CD4+ T cells in the development of autoimmune peripheral neuropathy. Nevertheless, much remains unknown about effector mechanisms, ontogeny, and peripheral nerve localization of PNS-reactive CD4+ T cells. In this study, we demonstrate that IL-21 production is a hallmark of pathologically expanded, clonally related CD4+ T cells in infiltrated peripheral nerves. Genetic IL-21R deficiency completely protects against neuropathy development, demonstrating that IL-21 signaling is required for autoimmune pathogenesis. Finally, we show that IL-21 upregulates the chemokine receptor CXCR6 in CD4+ T cells, suggesting a role for IL-21 in CD4+ T cell positioning. These findings point to IL-21/IL-21R and CXCR6/CXCL16 as promising targets for novel therapies in inflammatory neuropathies.

IL-21R is expressed by various immune cell types, which implies that IL-21 has broad cellular targets^40^. Previous studies have highlighted the role of IL-21 in CD8+ T cell or B cell activation and differentiation^27^. However, the frequency and absolute numbers of CD8+ T cells and B220+ B cells were unchanged in peripheral nerves of IL-21R deficient NOD.Aire^GW/+^ mice. Instead our data demonstrate lower numbers of CD4+ T cells, suggesting that IL-21 functions in an autocrine fashion to increase CD4+ T cells within peripheral nerve infiltrate. Of note, we have previously reported a pathogenic role for IFN-γ and IL-10-producing CD4+ T cells in SAPP^32,33^, and IL-21R deficiency results in a substantial decrease in these cells within peripheral nerves. Finally, we show that IL-21 signaling functions to upregulate CD4+ T cell expression of the chemokine CXCR6, suggesting a potential role for CXCR6 in pathogenic CD4+ T cell accumulation in peripheral nerves.

CXCR6-expressing T cells are well-studied in anti-cancer immunity, where CXCR6 is used as a marker of resident memory T cells (T_RM_)^42^. Within tumors, CXCR6 positions T cells next to perivascular dendritic cells that express the CXCR6 ligand, CXCL16^45^. At the same time, CXCR6-expressing T cells are enriched in inflamed tissue of patients with psoriasis and inflammatory arthritis,^46^ suggesting a pathogenic role for CXCR6 in these autoimmune diseases. Indeed, genetic CXCR6 deficiency is protective in mouse models of arthritis^47^, and antibody-mediated blockade of CXCR6 or CXCL16 ameliorates disease in a mouse model of multiple sclerosis^48,49^. Together, these findings demonstrate a critical role for CXCR6/CXCL16 interactions in these autoimmune conditions. Our data suggest that CXCR6/CXCL16 interactions are also critical in PNS autoimmunity. scRNAseq analysis of infiltrated peripheral nerves shows accumulation of CXCR6-expressing CD4+ T cells and demonstrate that CXCR6 overexpression increases accumulation of CD4+ T cell numbers within nerves. Moreover, high levels of CXCL16 are expressed by macrophages and dendritic cells in peripheral nerve infiltrates, suggesting that CXCR6/CXCL16 interactions promotes localization of pathogenic CD4+ T cells to peripheral nerves.

Within infiltrated peripheral nerves, we identified CD4+ T cells associated with an expanded clonotype that were capable of expressing multiple pathogenic cytokines (IL21, IFN-y, and IL-10). This same set of cytokines are also expressed by Senescence Associated T cells (SATs), a CD153-expressing CD4+ T cell population associated with aging and inflammation^34,50,51^. SATs have been proposed to underlie the increased risk of autoimmunity, since anti-CD153-mediated depletion of SATs ameliorated disease in a mouse model of lupus^50^. Whether multi-functional Tph-like cells in inflamed peripheral nerves also express cell senescence features and accumulate with age, however, remains to be explored. However, it is intriguing that the incidence of inflammatory neuropathies increases with advancing age, with the peak age of onset between 70-79 years for CIDP and >60 years for GBS^52,53^. These findings suggest that age is a predisposing factor in the development of inflammatory neuropathies.

Collectively, our findings reveal a number of potential new therapeutic targets for inflammatory neuropathies. First, therapies targeting IL-21/IL-21R are under development for Type 1 Diabetes, Rheumatoid Arthritis, and psoriasis^23^, and our findings suggest that therapies blocking IL-21 or IL-21 signaling may be effective as a novel therapeutic target for inflammatory neuropathies. Second, therapies that target immune cell localization have been effective in inflammatory colitis and other immune-mediated diseases, and our data suggest that blocking CXCR6/CXCL16 receptor/ligand interactions may be efficacious for mitigating T cell localization to the peripheral nerves. Finally, our finding in this study that the most differentiated, clonally expanded CD4+ T cells can be triple cytokine producers suggests that therapies that target multiple cytokine signaling pathways, such as JAK/STAT inhibitors, may be a new therapeutic strategy for inflammatory neuropathies.

## Material and Methods

### Sex as a Biological Variable

We have previously reported that neuropathy age-of-onset and incidence is higher in female NOD.Aire^GW/+^ mice compared to male^32^. Here, we use both female and male NOD.Aire^GW/+^ IL21R^KO^ mice to show that IL-21 signaling is required for neuropathy in both sexes (**Fig. 4A and Supp. Fig. 3A)**. We subsequently focus on female mice in other studies given the earlier onset and higher incidence in NOD.Aire^GW/+^ females.

### Mice

NOD.Aire^GW/+^ mice^9^ and NOD.Cg-Prkdc^scid^/J (NODSCID, JAX 001303) mice were housed in a pathogen-free (SPF) barrier facility at UCLA. NOD.Aire^GW/+^IL21R^KO^ mice were produced by crossing together NOD.129(Cg)-Il21r^tm1Wjl^/DcrMmjax (NOD.Il21RKO, JAX 050918) mice and NOD.Aire^GW/+^ mice. In all experiments, female mice were used. All experiments with mice were approved by the UCLA Animal Research Committee.

### Neuropathy assessment and nerve conduction studies

Neuropathy was determined as previously described^54^. EMGs were performed using a TECA Synergy N2 EMG machine as previously described^55^. Compound Muscle Action Potentials (CMAPs) were recorded following the stimulation of the sciatic nerve with 0.1ms duration, 1Hz frequency and 20mA intensity stimulus with low pass filter set to 20Hz and high pass to 10kHz.

### Histology and Immunostaining

The forelimb nerves were dissected for histology and immunostaining. Paraffin-embedded nerve samples were stained with H&E and used for semi-quantitative immune cell infiltrate scoring on a scale from 0 to 4 as previously described^10,32^. 3-4 non-overlapping microscopic fields were evaluated per nerve. For immunostaining of frozen nerve sections, tissues were fixed in 4% paraformaldehyde overnight and cryopreserved with sucrose 30% for 1-2 days. Nerve samples were embedded in OCT medium (FisherScientific). The frozen blocks were prepared by placing the embedding molds in ethanol cooled by dry ice. 10 μm sections were cut from the nerves and collected on Superfrost/Plus slides (Fischer Scientific). For immunostaining, slides were incubated with primary antibodies overnight at 4°C; the next day slides were washed, followed by application of secondary antibodies for 1 hour. Slides were mounted with Fluoromount-G with DAPI (Invitrogen, 00-4959-52). The list of the antibodies used for immunofluorescent staining can be found in Table 1. Images were acquired using a ZEISS Axiocam 208 or ZEISS Axiocam Observer and processed using ImageJ/Fiji software.

### Flow cytometric analysis

Following cardiac perfusion with PBS, single cells were isolated from the spleen, lymph nodes (2 brachial and 2 inguinal), and sciatic nerves as previously described^54^. Briefly, chopped sciatic nerves were digested with 1mg/ml collagenase IV and passed through a 20-gauge needle. Spleen, lymph node, and digested sciatic nerve samples were passed through 40µm filters and washed with PBS, yielding single cell suspensions.

For intracellular cytokine staining, cells were stimulated with PMA (50ng/ml), ionomycin (1ug/ml), BFA (1x) and Monensin (1x) for 4 hours. Cells were stained with antibodies against cell surface proteins, then fixed with Fix & Perm Medium A (Invitrogen, GAS001S100) followed by permeabilization with Fix & Perm Medium B (Invitrogen, GAS002S100) for cytokine staining. Antibodies used for flow cytometry are listed in Table 1. A BD Fortessa Cell Analyzer or Attune NxT Flow Cytometer was used to perform flow cytometry. The flow cytometry data were analyzed using FlowJo software v10.

### Cell sorting

Cells were sorted using a BD FACS Aria Cell Sorter at the UCLA BSCRC Flow Cytometry Core, mouse CD4 isolation kit (Miltenyi, 130-104-454), or mouse CD8 isolation kit (Miltenyi, 130-104-075).

### CXCR6 virus preparation and transduction

Cxcr6 expressing plasmid was generated by cloning *Cxcr6* gene block amplified from ORF (GenScript) into MSCV-IRES-GFP backbone (Addgene # 20672). Retrovirus was produced by co-transfecting Phoenix-ECO (American Type Culture Collection CRL-3214) with transfer plasmid, MSCV-Cxcr6-IRES-GFP or pMSCV-IRES-mCherry (Addgene # 52114) and pCL-Eco (Addgene #12371) using TransIT-293 transfection reagent (Mirus Bio Cat. 2705). Media was changed 16 hours post-transfection. Retrovirus was collected in the two following days and supernatant was stored at −80C for transduction. Primary murine CD4 T cells were stimulated overnight with platebound anti-CD3/anti-CD28 beads before transduction. Transduction was performed on Days 1 and 2 by spinoculation at 2000 x g for 90 minutes at 32°C using low acceleration and minimal deceleration. Transduced cells were incubated for 4 days with plate-bound anti-CD3/anti-CD28 in media supplemented with hIL-2 (0.0344 units/mL, Peprotech #200-02). Transduction efficiency was assessed for each experiment by flow cytometry.

### Single cell RNA sequencing and analysis

scRNA sequencing was performed for CD45+ isolated cells from the sciatic nerves of 4 neuropathic NOD.Aire^GW/+^ mice. We performed 10x Genomics Single Cell Immune Profiling [V(D)J + 5’ Gene Expression] to the UCLA Technology Center for Genomics and Bioinformatics (TCGB) core. Analysis was performed using the Seurat package (4.3.0) on R Studio (version 4.2.3 – “Shortstop Beagle”). Raw sequences were processed and mapped to reference genome (mm10) using Cell Ranger. Outputs of the Cell Ranger pipeline were read into R Studio with the Read10X function, generating single cell-level transcript counts of each gene. Cells with fewer than 300 RNA count, greater than 5000 features, and greater than 20% mitochondrial gene were excluded from the analysis. Data were normalized and log-transformed with the NormalizeData function, and variable features were identified using the FindVariableFeatures function. To maximize cell numbers, this dataset was integrated with a previously published dataset from 3 neuropathic NOD.Aire^GW/+^ mice (GSE 180498)^56^. Integration anchors were identified amongst the Seurat object inputs. A single integrated analysis was performed on all *Cd4*-expressing cells. The standard workflow for visualizing and clustering occurred; data was scaled with ScaleData and linear dimensional reduction (RunPCA) was performed. Cell clusters were determined by the FindClusters function and visualized by Uniform Manifold Approximation and Projection (UMAP). Cell types were identified by expression of canonical markers described in existing literature and within the Immgen RNASeq Skyline database^57^. Differentially expressed genes (DEGs) which were conserved across datasets were identified with FindConservedMarkers.

The Slingshot (2.6.0) and SingleCellExperiment (1.20.1) R packages were used to characterize global structure and predict lineages based on cluster relationships. Slingshot performed trajectory inference with the dimensionality reduction produced by PCA and a set of cluster labels. To perform clonotype analysis, the filtered_contig_annotations.csv output from 10X Genomics Cell Ranger was loaded from each VDJ alignment folder to generate the dataset used in scRepertoire (1.7.0). Extraneous prefixes of cell barcodes were removed with the stripBarcode function. A single list object of TCR genes and CDR3 sequences by cell barcode was combined with the integrated CD4+ T cell Seurat object using the combineExpression function. The different clonotypes frequencies were projected onto the Seurat object’s UMAP. CellChat R package (version 0.5.5) was used to make inferences about potential cell-cell Interactions, as previously described^56^.

### Statistical analysis

Statistical analysis was performed using GraphPad Prism 9 or R for scRNA sequencing analysis. Unpaired t tests were used to compare two groups while paired t tests were used for matched samples. Mann-Whitney tests were used to compare two groups with non-parametric distribution. One-way ANOVA with Bonferroni post-test analyses were used for the comparison of multiple groups. Data with more than one variable was compared using a two-way ANOVA followed by Bonferroni’s multiple comparisons test. For neuropathy incidence curves, a log-rank (Mantel-Cox) test was used. Bonferroni’s adjusted p-values were reported for differentially expressed genes and Benjamini-Hochberg adjusted p-values were used in Pathway Analysis. Fold changes were calculated as (B-A)/A. An adjusted p-value of less than 0.05 was considered significant. Bargraph and dot plot data are presented as mean±SEM with the following symbols: *p<0.05, **p<0.01, *** p<0.001.

## Study approval

All experiments with mice were approved by the UCLA Animal Research Committee.

## Data availability

The datasets for scRNAseq and scTCRseq generated during the current study are available in the Gene Expression Omnibus (GEO) repository under accession numbers GSE252646 and GSE252647 (https://www.ncbi.nlm.nih.gov/geo/).

## Author contributions

MS, JKW, MGL and MAS designed research studies. MS, MB, MM conducted experiments, and acquired data. MS, MB, EM, SZ, HCC, MM, WH analyzed data. JKW and SZ provided reagents. MS, MB, EM, MM, MAS wrote the manuscript.

## Acknowledgements

MAS is supported by the NIH (NS107851, AI143894, DK119445), GBS/CIDP Foundation International, and National Organization of Rare Diseases. MAS and JKW is supported by DOD (USAMRAA PR200530).

## Supplemental Figure Legends

**Supp. Fig. 1. (A)** Heatmap showing top 14 DEGs for each cluster with key identifying genes bolded in left. **(B)** Feature plots of CD4+ T cells showing expression of regulatory T cell (Treg) genes (*Foxp3*, *Il2ra).* **(C)** Published UMAP (GSE 180498) of CD45+ T cells integrated from B6.WT, NOD.WT or NOD.Aire^GW/+^ sciatic nerves. **(D)** Feature plots of CD45+ T cells showing expression of *Il21,* split by genotype (NOD.WT and NOD.Aire^GW/+^).

**Supp. Fig. 2. IL-21 producing Tph-like cells in sciatic nerves of neuropathic NOD.Aire^GW/+^ mice. (A)** Representative flow cytometry plot of intracellular IL-21 staining of peripheral nerve CD4+ T cells from NOD.WT vs. neuropathic NOD.Aire^GW/+^ sciatic nerves. **(B)** Gating strategy for flow cytometric analysis of Tfh (CD4+ICOS+PD1+CXCR5+) and Tph-like cells (CD4+ICOS+PD1+CXCR5-CXCR6+). **(C)** UMAP plot of CD4+ T cells with overlaid Slingshot pseudotime trajectories.

**Supp. Fig. 3. IL-21 signaling upregulates T cell cytokine and chemokine expression. (A)** - Neuropathy free incidence curve for male NOD.Aire^GW/+^ IL-21R^KO^ vs. NOD.Aire^GW/+^ IL-21R^Het^ and NOD.Aire^GW/+^ IL-21R^WT^ mice. Mantel-Cox test; **p<0.01. **(B)** Flow cytometric analysis of intracellular IL-21, IL-10, and IFN-y staining of CD4+ T cells in peripheral nerves of Aire^GW/+^ IL-21R^WT^ vs. NOD.Aire^GW/+^IL-21R^KO^ mice. **(C)** *Cxcr6* expression levels after IL-21 stimulation of CD4+ T cells at different timepoints. Data is plotted from Affymetrix gene expression data reported in Table S3 of Kwon et al^58^. The two curves represent the two microarray probes that map to CXCR6 (probe IDs 1425832_a_at and 1422812_at). Each point is an average of 5 biologic replicates.

**Supp. Fig. 4. CXCR6 and CXCL16 expression is upregulated in neuropathic NOD.AireGW/+ peripheral nerves. (A)** Feature plots of CD45+ T cells showing expression of *Il21,* split by genotype (NOD.WT and NOD.Aire^GW/+^). UMAP is from Supp. Fig. 1C.

---

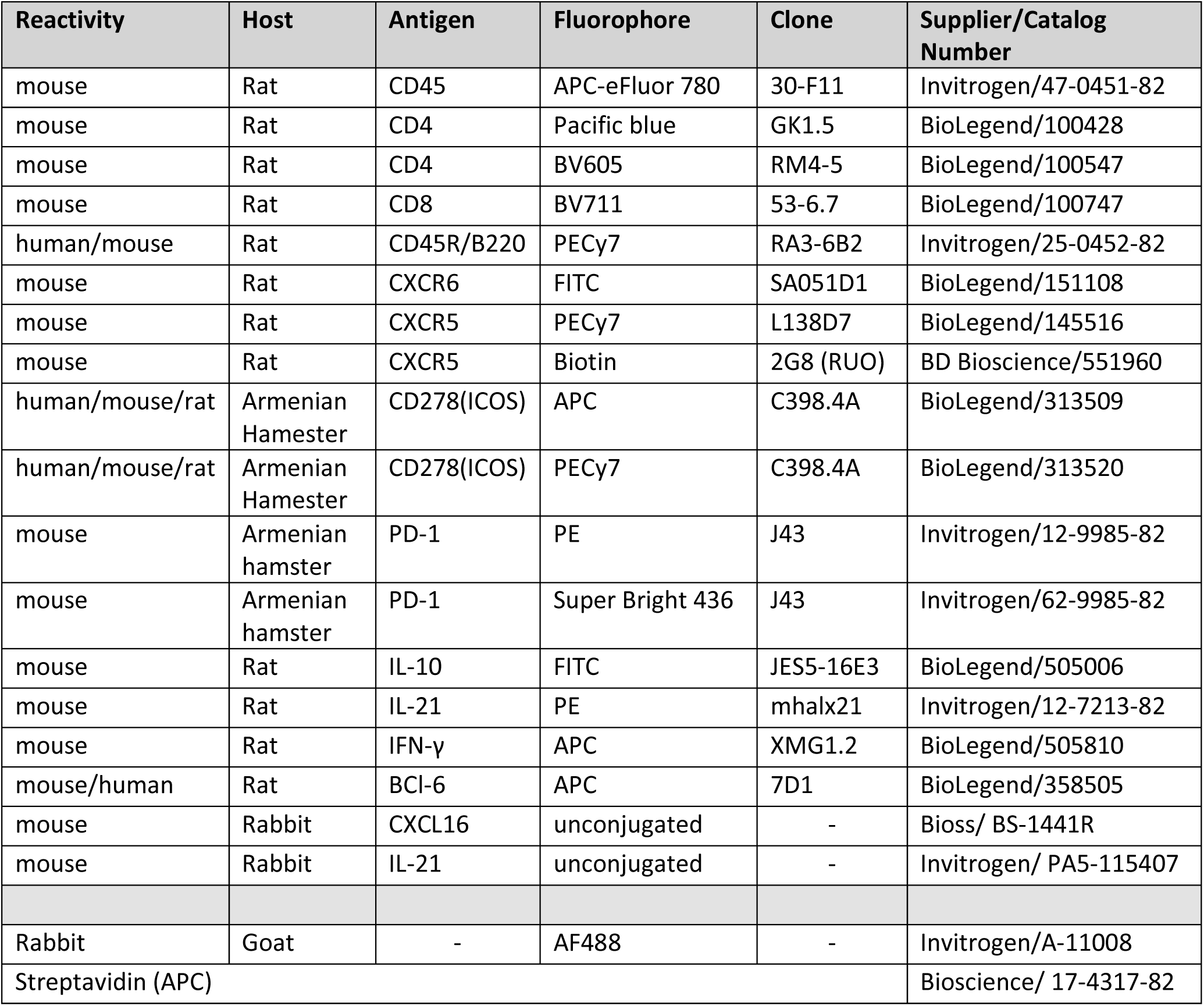

## Notes

Conflict of Interest Statement: The authors have declared that no conflict of interest exists.

### Competing Interest Statement

The authors have declared no competing interest.

