## Supplementary figures and images for "A pathologically expanded, clonal lineage of IL-21 producing CD4+ T cells drives Inflammatory neuropathy"

### Supplemental Figure1

##### Supplemental Figure 1

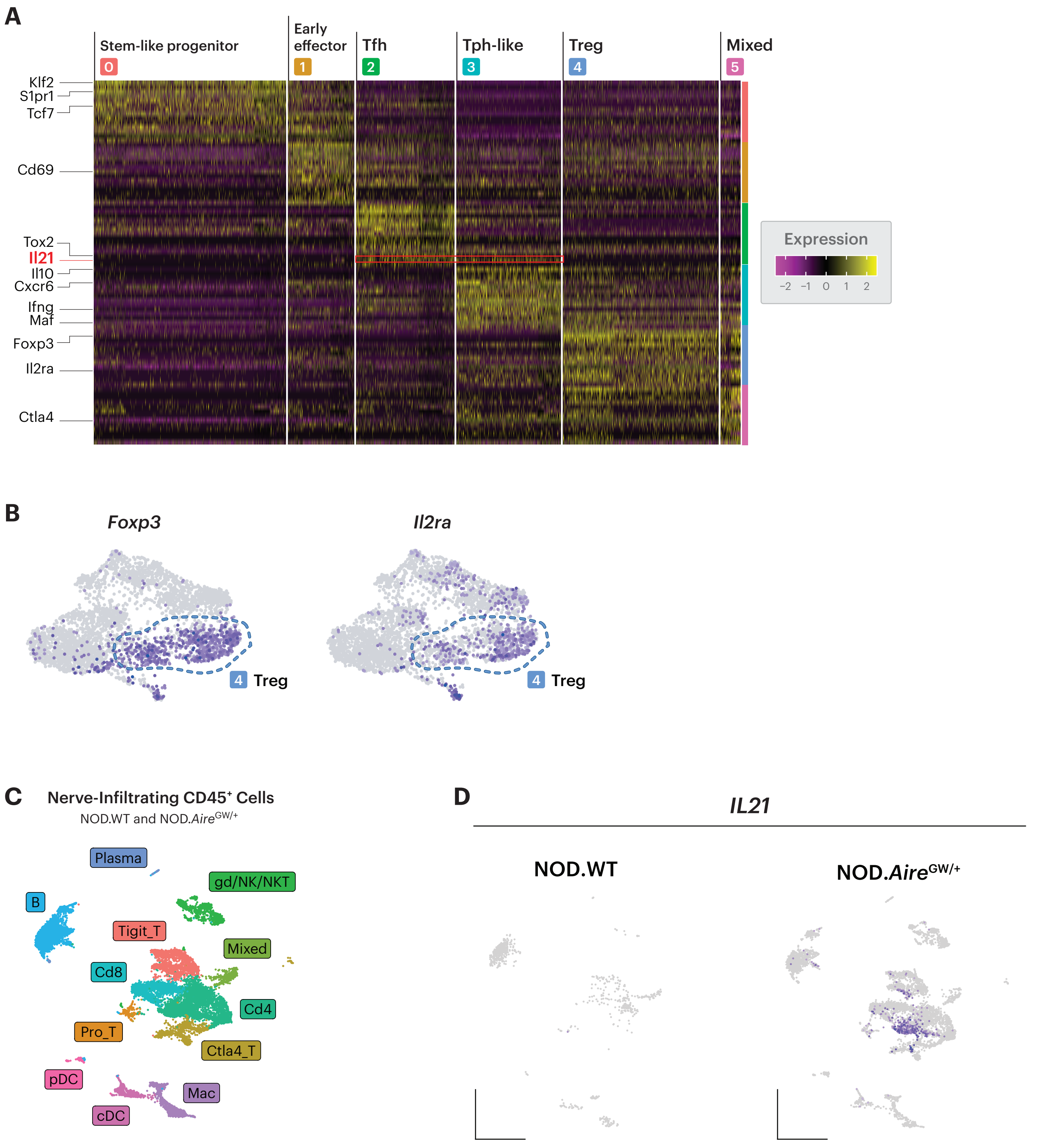

### Supplemental Figure 2

#### Supplemental Figure 2

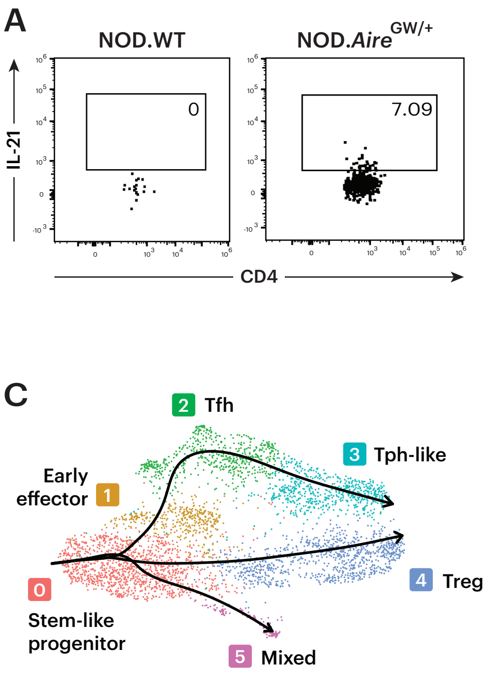

### Supplemental Figure 3

##### Supplemental Figure 3

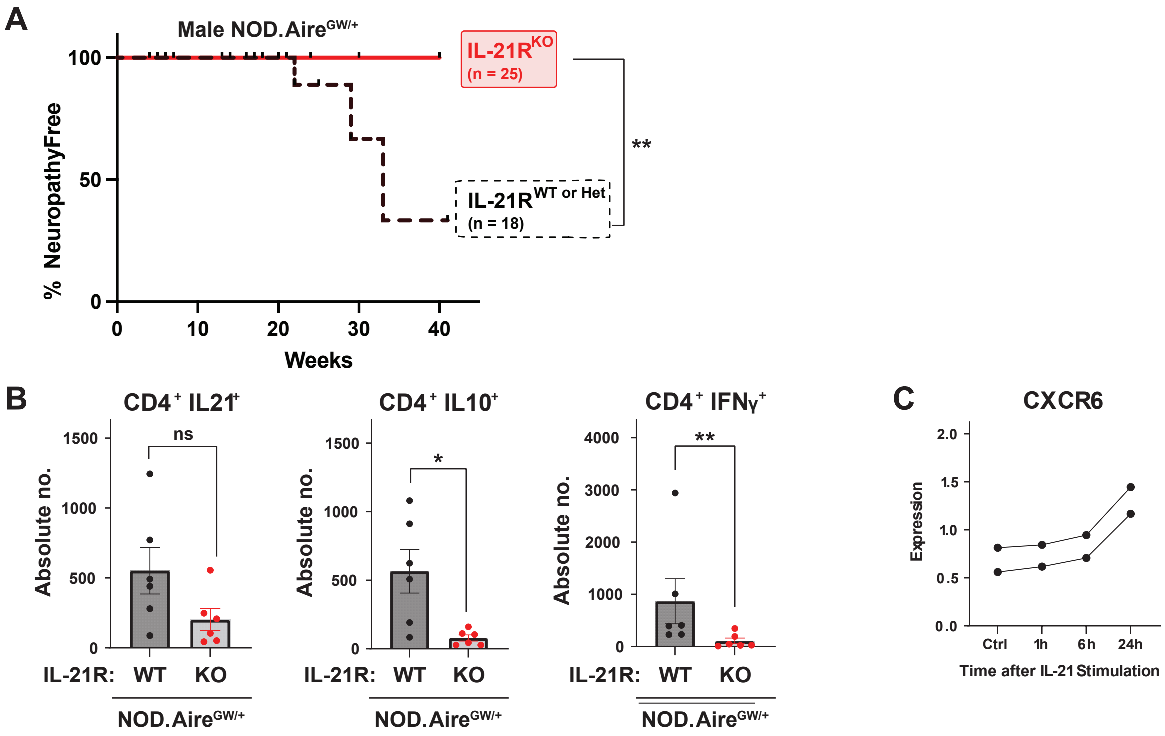

### Supplemental Figure 4

##### Supplemental Figure 4

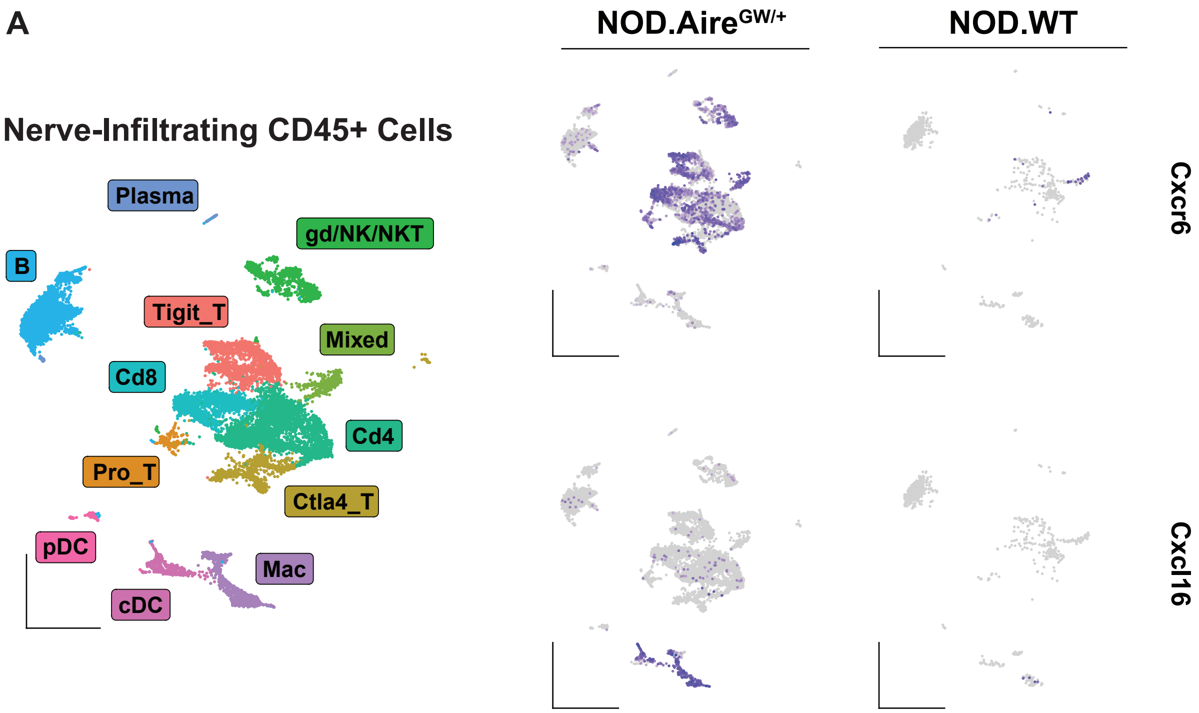
